# The decline of malaria in Vietnam, 1991-2014

**DOI:** 10.1101/151456

**Authors:** Sandra M Goldlust, Phung Duc Thuan, Dang Duy Hoang Giang, Ngo Duc Thang, Guy E Thwaites, Jeremy Farrar, Ngo Viet Thanh, Tran Dang Nguyen, Bryan T Grenfell, Maciej F Boni, Tran Tinh Hien

## Abstract

A central component of malaria control initiatives throughout the world is the use of artemisinin-based combination therapies (ACTs) for treatment of uncomplicated *P. falciparium* malaria. Despite the well-documented clinical efficacy of ACTs, the population-level effects of ACT case management on malaria transmission have not been studied thoroughly until recently. An ideal case study for the population-level effects of artemisinin use can be found in Vietnam, where a major increase of malaria cases in the 1980s was followed by the gradual adoption of artemisinin-based clinical case management. We assembled annual data from Vietnam’s National Institutes for Malariology, Parasitology, and Entomology showing the degree to which artemisinin therapies were adopted in different provinces, the effort placed on vector control, and the funding available to provincial malaria control programs, from 1991 to 2014. Data on urbanization were also collected for this period. We found that a 10% increase in the artemisinin proportion of treatments procured by a provincial control program corresponded to a 32.8% (95% CI: 27.7 – 37.5%) decline in estimated malaria cases; the association persisted and the effect size was nearly unchanged if confirmed cases or suspected cases were used. There was no consistent effect of vector control on malaria cases in Vietnam as a whole, nor was any effect found when the data were broken up regionally. The association between urbanization and malaria was generally negative and sometimes statistically significant. This was most pronounced in the central region of Vietnam, where a 10% increase in urbanization corresponded to a 43.3% (95% CI: 21.6 – 58.9%) decrease in suspected malaria incidence; this association was not statistically significant if confirmed cases or estimated cases were used. The decline of malaria in Vietnam from 1991 to 2014 can largely be attributed to the rapid adoption of artemisinin-based drugs. Recent analyses of aggregated data from Africa have shown that insecticide-treated nets have had the greatest effect on lowering malaria prevalence over the past fifteen years, suggesting that the success of different types of malaria interventions is region specific. Continuing global efforts on malaria elimination should focus on both vector control measures and increased access to artemisinin-combination therapies.

## Introduction

Over the past fifteen years, scale-up in key tools to prevent and treat malaria has contributed to a dramatic reduction in transmission worldwide [1]. Principal among these tools have been insecticide-treated nets, indoor residual spraying of insecticide, and artemisinin-based combination therapies (ACTs) – the most effective antimalarial therapies currently available for treatment of uncomplicated *P. falciparum.* In 2005, the World Health Organization began recommending ACTs as the first-line therapy for uncomplicated *P. falciparum* [2], although treatment with artemisinin-containing antimalarials had already begun to be put into place in areas confronted with parasite resistance to chloroquine, sulfadoxine-pyrimethamine, and mefloquine [3–5]. Early clinical studies on artemisinin derivatives in the 1990s [6–10] and larger trials of ACTs in the 2000s [11–15] demonstrated high clinical efficacy and a rapid killing rate, which would later make ACTs the first choice of national anti-malaria programs in malaria-endemic countries.

Beyond its well-established clinical efficacy, artemisinin may have population-level benefits for malaria control due to its effect of reducing post-treatment carriage of gametocytes [16–19] - the sexual stage of malaria transmitted from human peripheral blood to *Anopheles* mosquitoes. Treatment of *P. falciparum* with an artemisinin-containing antimalarial results in the rapid killing of parasite asexual stages (99% daily kill rate [20, 21]) and, when combined with a partner drug, results in undetectable parasitaemia by microscopy after three days of treatment [10, 22]. Low parasite densities in the blood generally indicate that patients are less likely to transmit the sexual stages of the parasite to mosquitoes [23, 24]. However, the long-term population-level effects of ACT case management on parasite transmission are only now beginning to be documented [1, 25–27]. The fact that ACTs are typically introduced as a component of comprehensive malaria control efforts makes it challenging to isolate the effectiveness of ACTs from that of other concurrently introduced control strategies, such as indoor residual spraying (IRS) and utilization of insecticide-treated bed nets (ITNs).

One of the best countries to use as a case study for the long-term effects of artemisinin use in malaria case management is Vietnam. Following a major epidemic of chloroquine-resistant *P. falciparum* in the late 1980s, Vietnam implemented a new national malaria control program into which it introduced case management with artemisinin-containing antimalarials as a possible solution to the high malaria incidence the country was experiencing at the time; in addition, vector control measures were introduced and health capacity was strengthened. The incidence of malaria in Vietnam subsequently declined [5, 28]. Malaria incidence, antimalarial use, vector control efforts, and various measures of health systems strengthening were recorded in annual reports produced by Vietnam’s National Institutes for Malariology, Parasitology, and Entomology (NIMPE). An analysis of these data for the provinces in the southern part of the country from 1991-2010 showed that the strongest association with reduced malaria incidence was the proportion of stocked or ordered antimalarial drugs that were artemisinin derivatives [27]. In the present study, we co-analyze these data with national reports collected for Vietnam’s central and northern provinces, extending the analysis through 2014. We make several new observations about the factors that are most closely associated with reduced malaria incidence and about the regional variation in associations in the northern, central, and southern parts of the country. We also consider the case reporting process and diagnostic capabilities of Vietnam’s national malaria program in order to improve our estimates of the malaria case burden in Vietnam over the past two and a half decades. Our findings underscore the robustness of the association between adoption of artemisinin-containing antimalarials for case management of *P. falciparum* and malaria decline, and we hope they will inform the development of effective public health strategies for eliminating malaria in Vietnam in the near future.

## Results

### Changes in malaria transmission

All measures of malaria incidence in Vietnam declined significantly between 1991 and 2014, with the majority of the decline occurring in the 1990s (Figure 1). The total number of suspected malaria cases in Vietnam declined from 1,290,250 cases in 1992 to 27,868 cases in 2014, corresponding to a 98.3% reduction in the incidence of suspected cases. The incidence of confirmed cases decreased by 94.9% over this time period, with the number of confirmed cases declining from 224,923 in 1992 to 14,941 in 2014. The numbers of severe malaria cases declined from 24,022 in 1992 to 65 in 2014 and the number of deaths declined from 2,702 in 1992 to 9 in 2014. Similarly, the number of estimated malaria cases declined between 1992 and 2014 from 586,172 to 17,939. The declining trend in incidence between 1991-2014 was consistent across all provinces in Vietnam (Figure 2).

**Figure 1.**
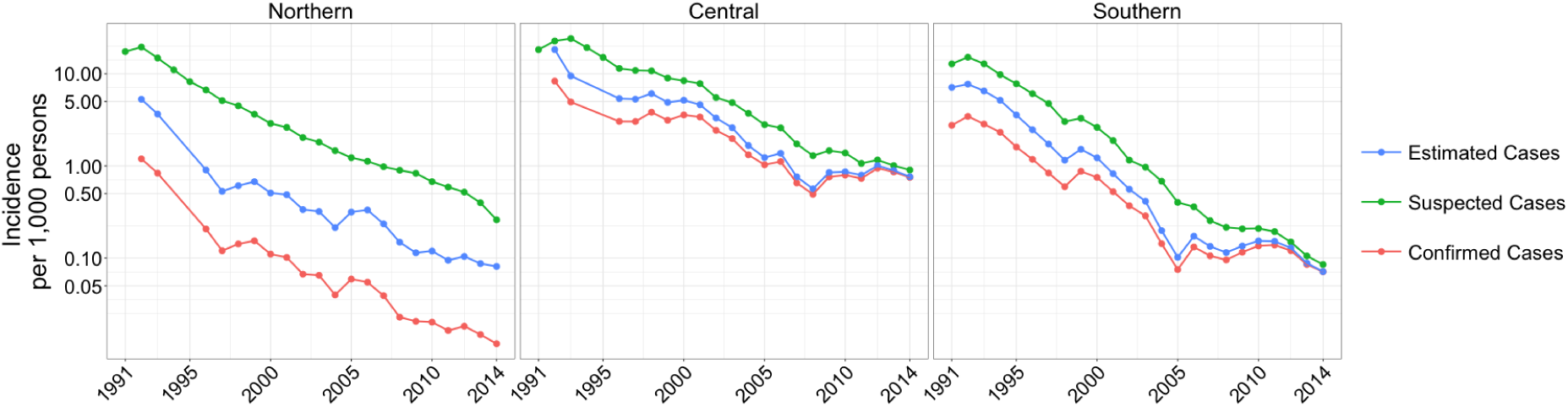
Incidence of suspected, confirmed, and estimated cases of malaria per 1,000 person-years by region from 1991-2014 on a log-transformed scale where labels correspond to raw (unlogged) values. In 2014, there were 473 confirmed cases in the northern region, 12,006 confirmed cases in the central region, and 2,462 cases in the southern region.

**Figure 2.**
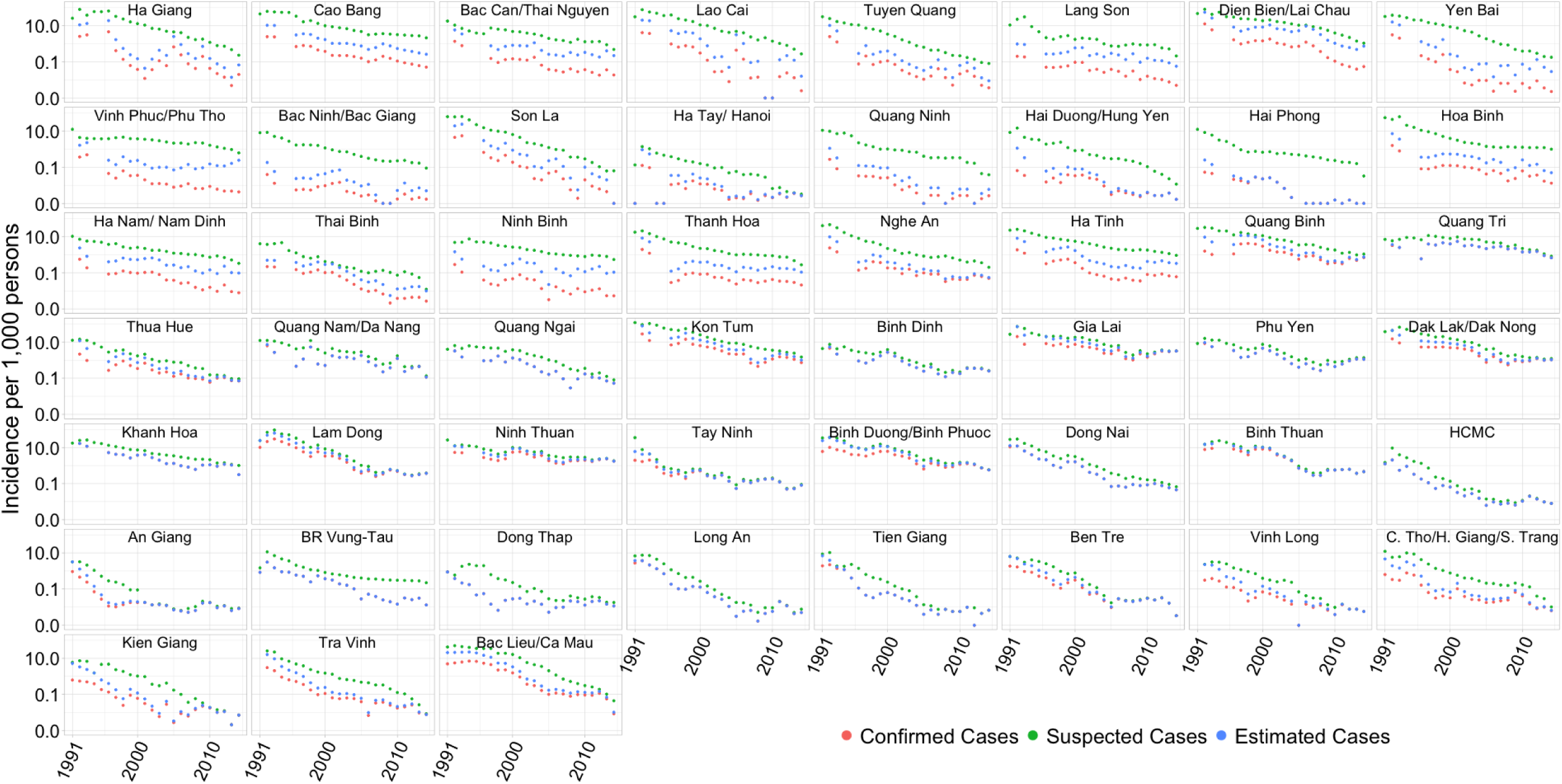
Incidence of suspected, confirmed, and estimated cases of malaria, by province (1992-2014). Provinces are arranged approximately by decreasing latitude (north to south) from top to bottom, and left to right. The *y*-axis is log-transformed, but the labels correspond to raw (unlogged) values and the “0.0” label on the *y*-axis corresponds to true zero.

### Malaria burden in Vietnam, 2014

Table 1 reports the number and incidence of malaria cases in 2014 for the 22 provinces (arranged approximately from the northern to the southern region) that reported more than 50 suspected cases of malaria. In 2014, malaria transmission was greatest in Gia Lai province, which had an estimated 4,386 cases and an annual incidence of 3.2 estimated cases per 1,000 individuals. Using our estimation of malaria cases, we found malaria transmission in 2014 was most heavily concentrated in the central region of Vietnam, and that incidence varied greatly among the northern provinces (Figure 3A). Mapping the positive predictive value of clinical malaria diagnosis, *q,* in 2014 revealed that *q* tended to be much greater in the south compared to the north (Figure 3B). The difference in positive predictive value between the northern and southern regions can be attributed to northern provinces having high numbers of suspected malaria cases while reporting very few slide-confirmed cases, which resulted in low positive predictive values in the north. For the years 2010-2014, the average positive predictive values ranged from 0.10% in Hai Phong (located in the northern region) to 99.2% in Binh Thuan (located in the southern region). Excluding the northern provinces (those north of and including Ha Tinh) and an outlier in Ba Ria-Vung Tau, the average positive predictive value ranged from 31.4% in Hau Giang/Can Tho/Soc Trang to 99.2% in Binh Thuan province; this is the expected range for the positive predictive value of malaria clinical diagnosis according to senior IMPE/NIMPE staff.

**Figure 3.**
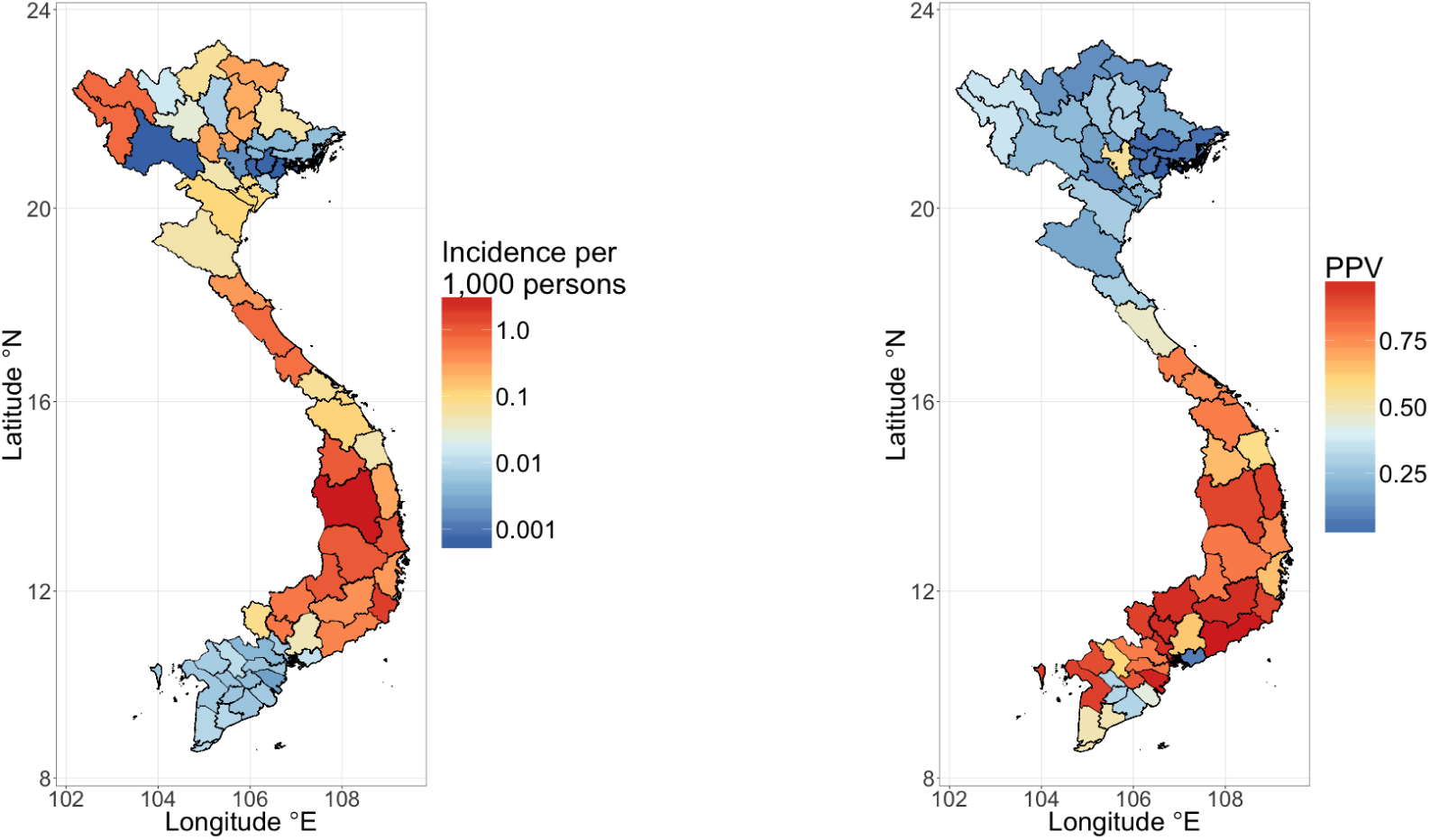
**A)** Incidence of malaria in Vietnam, 2014. Incidence is calculated per 1,000 person-years using the estimated number of cases. **B)** Average positive predictive value, *q,* by province for the years 2010-2014.

**Table 1.** Malaria cases in 2014, by province

| Province Name (Latitude °N) | Estimated Cases | Suspected Cases | Confirmed Cases | Incidence of<br>Estimated Cases | Incidence of<br>Suspected Cases | Incidence of<br>Confirmed Cases |
| --- | --- | --- | --- | --- | --- | --- |
| Dien Bien/Lai Chau (21.8° / 22.4°) | 700 | 1022 | 51 | 0.734 | 1.072 | 0.053 |
| Thanh Hoa (20.1°) | 374 | 957 | 69 | 0.107 | 0.274 | 0.020 |
| Nghe An (19.2°) | 163 | 604 | 144 | 0.054 | 0.199 | 0.047 |
| Ha Tinh (18.3°) | 408 | 1146 | 74 | 0.325 | 0.913 | 0.059 |
| Quang Binh (17.5°) | 647 | 930 | 596 | 0.745 | 1.071 | 0.686 |
| Quang Tri (16.8°) | 413 | 502 | 413 | 0.670 | 0.814 | 0.670 |
| Thua Hue (16.5°) | 80 | 103 | 78 | 0.071 | 0.091 | 0.069 |
| Quang Nam/Da Nang (15.5° / 16.0°) | 283 | 331 | 283 | 0.114 | 0.133 | 0.114 |
| Quang Ngai (15.1°) | 64 | 99 | 64 | 0.052 | 0.080 | 0.052 |
| Kon Tum (14.7°) | 485 | 716 | 350 | 1.002 | 1.479 | 0.723 |
| Binh Dinh (13.8°) | 376 | 395 | 376 | 0.248 | 0.261 | 0.248 |
| Gia Lai (13.8°) | 4386 | 4424 | 4367 | 3.183 | 3.211 | 3.170 |
| Phu Yen (13.1°) | 983 | 1199 | 983 | 1.108 | 1.351 | 1.108 |
| Dak Lak/Dak Nong (12.7° / 12.3°) | 2528 | 2921 | 2528 | 1.051 | 1.215 | 1.051 |
| Khanh Hoa (12.3°) | 376 | 1214 | 376 | 0.314 | 1.014 | 0.314 |
| Lam Dong (11.9°) | 465 | 479 | 465 | 0.369 | 0.38 | 0.369 |
| Ninh Thuan (11.7°) | 1033 | 1079 | 1033 | 1.750 | 1.828 | 1.750 |
| Binh Duong/Binh Phuoc (11.3° / 11.8°) | 1600 | 1669 | 1600 | 0.570 | 0.595 | 0.570 |
| Tay Ninh (11.4°) | 90 | 98 | 90 | 0.082 | 0.089 | 0.082 |
| Dong Nai (11.1°) | 127 | 188 | 127 | 0.045 | 0.066 | 0.045 |
| Binh Thuan (11.0°) | 559 | 559 | 559 | 0.463 | 0.463 | 0.463 |
| Ho Chi Minh City (10.8°) | 57 | 57 | 57 | 0.007 | 0.007 | 0.007 |

*Note.* Incidence measured per 1,000 persons. Provinces ordered by decreasing latitude (°N) from north to south. Only the 22 provinces with > 50 suspected cases in 2014 are shown.

### Malaria control measures between 1991-2014

Across Vietnam, the percentage of treatment courses ordered for *P. falciparum* malaria that contained artemisinin increased from 12.1% in 1992 to 92.9% in 2014. In 49 provinces, Spearman’s rank correlation tests revealed a significant positive correlation between time and the proportion of treatments for *P. falciparum* malaria containing artemisinin (*p* < 0.0001 in 48 provinces, and 0.0001 < *p* < 0.001 in Vinh Long). Using the Bonferroni correction, we considered Spearman p-values above 0.001 not to be statistically significant due to the large number of tests performed. The remaining two provinces (An Giang and Quang Ninh) showed an increasing trend in the proportion of artemisinin purchases through time, but the Spearman p-values for these trends were between 0.001 and 0.01 and thus do not guarantee statistical significance. The proportion of the population in Vietnam living in urban areas increased from 21.2% in 1995 to 33.1% in 2014. This trend was statistically significant in 42 provinces (*p* < 0.0001 in 38 provinces, and 0.0001 < *p* < 0.001 in 4 provinces). For the remaining 9 provinces, the trends showed increasing urbanization through time, but they were not statistically significant. As urbanization and artemisinin usage increased through time, we expected these covariates to be associated with the decline in malaria.

Unlike urbanization and the adoption of artemisinin, vector control measures did not show clear temporal trends when looking across the provinces. In Vietnam nationwide, the proportion of the population protected by vector control measures increased year-to-year from 1992-1997 (8.0% in 1992, 16.6% in 1996, 18.1% in 1997, 18.0% in 1998), and dropped to 9.7% in 2012 and 4.1% in 2014. In ten provinces, significant downward trends in vector control through time were detected (*p* < 0.0001 in 4 provinces, and 0.0001 < *p* < 0.001 in 6 provinces). The remaining provinces did not have strong support for either an upward or downward trend in vector control. The capacity of the health system, as measured by staff trainings per 100 persons was not strongly associated with time in the majority of provinces. The discretionary budget per capita decreased significantly over time in 24 provinces (*p* < 0.0001 in 17 provinces, and 0.0001 < *p* < 0.001 in 7 provinces), indicating that a budget increase was likely not a contributing factor to the decline of malaria in these provinces. However, it is important to remember that budgets and staff were reduced in many provinces because of the success of the control program; hence, this is a covariate whose estimates are difficult to interpret. No other trends were strongly supported in the remaining 27 provinces.

### Regression Results

In the nationwide analysis using all 51 provinces, after controlling for the proportion of the population living in urban areas and for the proportion of the population protected by ITNs or IRS, the proportion of treatments for *P. falciparum* that contained artemisinin was significantly (*p* < 0.001) and inversely associated with all three measures of malaria incidence (Figure 4). A 10% increase in the proportion of treatments containing artemisinin was found to be associated with a 32.8% (95% CI: 27.7 - 37.5%) reduction in the incidence of estimated cases. The proportion of the population living in urban areas was found to be significantly inversely associated with the number of suspected malaria cases (*p* < 0.001) and the number of estimated cases (*p* < 0.05), but not with the number of confirmed malaria cases (*p* = 0.48). The proportion of the population protected by vector control measures was not found to be significantly associated with any of the three measures of malaria incidence in the nationwide analysis.

When we additionally controlled for changes in health system capacity using the number of staff trainings per capita and the discretionary budget per capita, we similarly found that the proportion of treatments for *P. falciparum* containing artemisinin was significantly (*p* < 0.001) inversely associated with malaria incidence, as measured by suspected, confirmed, and estimated cases (Figure 5). Again, no significant associations were found between the proportion of the population protected by vector control and any of the three measures of malaria incidence. The proportion of the population living in urban areas was significantly inversely associated with the incidence of suspected cases (*p* = 0.004), but no associations were found between urbanization and estimated cases (*p* = 0.15) or confirmed cases (*p* = 0.29). The discretionary budget per capita was found to be positively but weakly associated with the number of suspected cases, with a 10% budget increase corresponding to a 1.35% (95% CI: 0.09 - 2.6%) increase in the number of suspected cases (*p* = 0.035). The number of staff trainings was not found to be significantly associated with any incidence measure.

**Figure 4.**
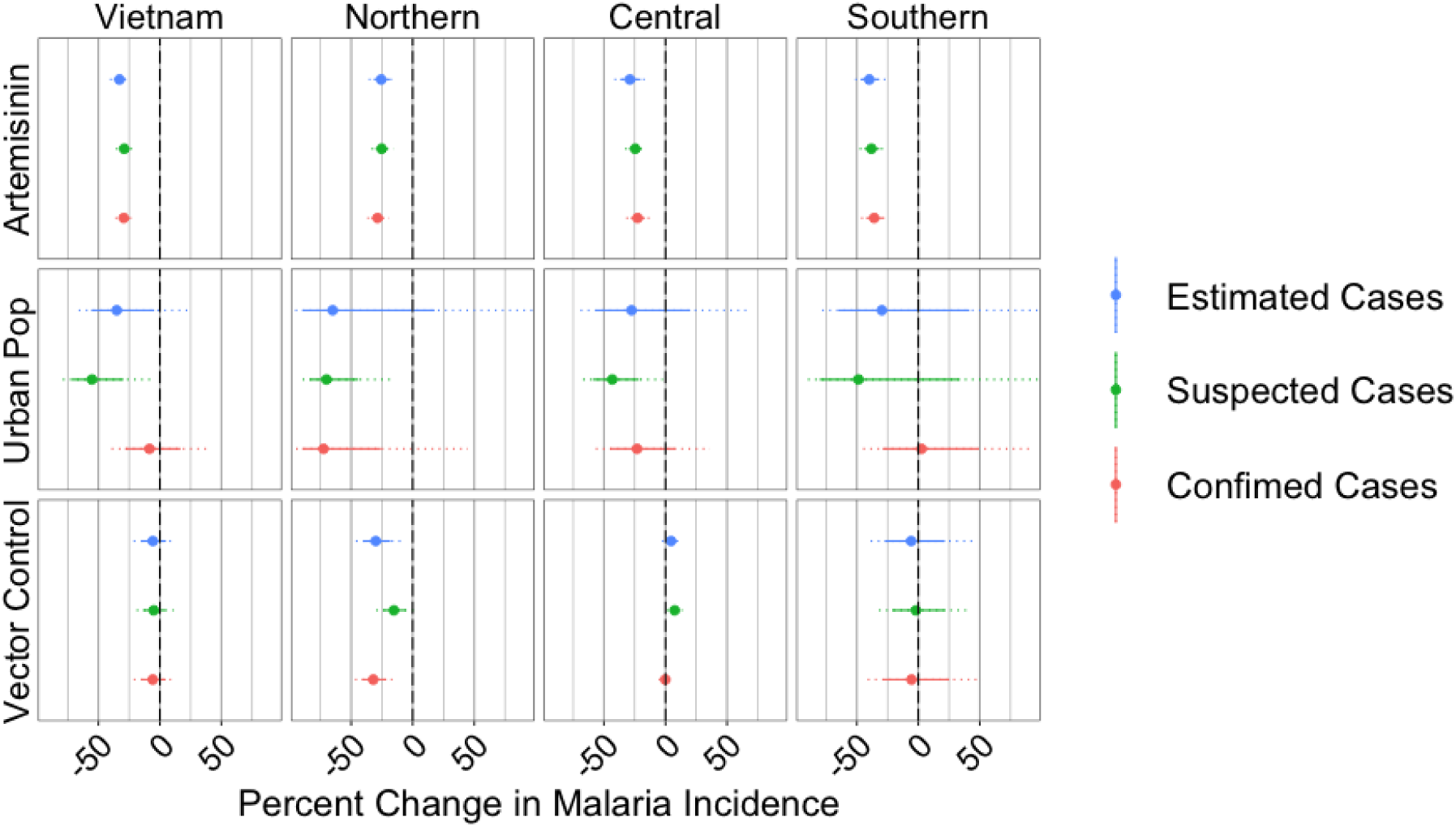
Percent change in malaria incidence associated with a 10% increase in the proportion of treatments for *P. falciparum* containing artemisinin (top row), proportion of the population living in urban areas (middle row), and proportion of the population protected by vector control measures (bottom row), by region and nationwide, as predicted by models using these three covariates only. The circle shows the mean effect size, the solid line shows the 95% confidence interval, and the dotted lines shows the 99.9% confidence interval. Outcome variable used in model is indicated by color. For clarity, the *x*-axis has been limited to range from -90% to 90%.

**Figure 5.**
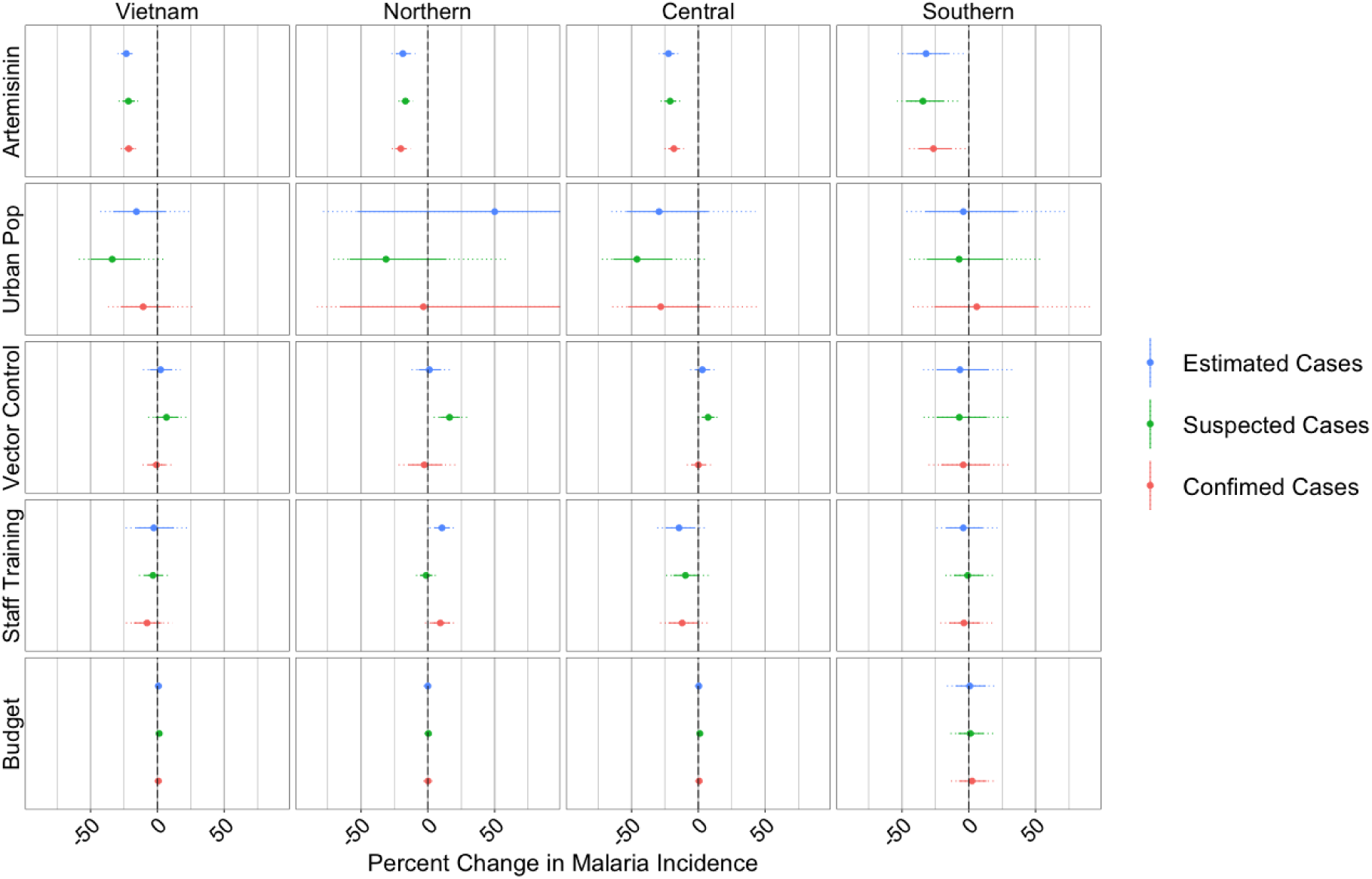
As in Figure 4, with the addition of covariates measuring health system capacity.

### Regional Variation in Regression Results

We found the proportion of purchased *P. falciparum* treatments that contained artemisinin was significantly (*p* < 0.001) inversely associated with all three measures of malaria incidence in all three regions (northern, central, southern) of the country, both with (Figure 4) and without (Figure 5) the inclusion of the health system capacity variables. A possible inverse association between the proportion of the population living in urban areas and malaria incidence can be seen for the central provinces, although it is only significant when looking at suspected malaria cases, suggesting that urbanization may have generally decreased incidence of non-specific rural febrile diseases (including malaria). The relationship between urbanization and malaria in the northern provinces is difficult to assess as the inferred associations were not robust across models and covariates. There was no evidence of a relationship between urbanization and malaria in the southern provinces.

Although the proportion of the population protected by vector control measures was not found to be significantly associated with malaria incidence when all provinces where analyzed together, a few significant associations were identified when the analysis was stratified by region. In the three-covariate models, the proportion protected by vector control measures was significantly inversely associated with all three measures of malaria incidence (*p* < 0.01) in the northern provinces. However, in the five-covariate models, this association only remained significant for incidence measured by suspected cases and the directionality of the effect was reversed.

In the three-covariate models for the central provinces, the proportion of the population protected by vector control measures was significantly positively associated with the number of suspected cases (*p* < 0.001) and with the number of estimated cases (*p* < 0.05). In the five-covariate models for the central provinces, the proportion of the population protected by vector control measures remained significantly positively associated with the number of suspected cases (*p* < 0.01), but was no longer significantly associated with the number of estimated cases. Neither the three-covariate model nor the five-covariate model revealed any significant associations between the population protected by vector control and any measure of incidence in the southern provinces. The lack of robustness in these results and the presence of positive associations between vector control and malaria led us to believe that the NIMPE/IMPE data did not contain any evidence for an effect of vector control on reducing malaria incidence.

The discretionary budget was not significantly associated with any of the three measures of incidence in any of the three regions (but see Appendix Figure S2a for results using alternative dose-to-course conversions). Staff trainings were significantly negatively associated with confirmed and estimated (*p* < 0.05) cases in the central region; however, in the northern region, staff trainings were significantly positively associated with confirmed (*p* < 0.01) and estimated (*p* < 0.001) cases. Overall, the health capacity variables do not have robust associations with malaria incidence; again, it is important to remember that as malaria incidence drops, budgets and staff numbers for malaria control programs will also be reduced. The results of the entire analysis were robust to method of calculation for the proportion of the treatments for *P. falciparium* containing artemisinin (Appendix Figures S2b and S2a).

## Discussion

The most robust statistical association detectable in the data reported by Vietnam’s National Institutes for Malariology, Parasitology, and Entomology from 1991-2014 is the negative association between malaria incidence – measured in three different ways as suspected cases, confirmed cases, and estimated cases – and the purchases of artemisinin-based drugs by provincial malaria control programs. Although the data only show that artemisinin-derivatives were purchased, every other description of the health system and antimalarial usage in the 1990s indicates that artemisinin drugs were used and even favored to treat *P. falciparum* malaria during this time because of their perceived efficacy [5, 28]. The significant negative association between the proportion of treatments for *P. falciparum* containing artemisinin and malaria incidence persists whether or not health system variables are included in the analysis and whether the data are broken down regionally or analyzed nationally. These findings are consistent with those of Peak et al [27] in their analysis of southern Vietnam from 1991-2010.

These results differ substantially from a recent continent-wide analysis in Africa showing that approximately 68% of the decline in malaria from 2000 to 2015 can be attributed to the use of insecticide-treated nets [1]. One reason may be the differential effort in Vietnam placed on ensuring access to ACTs versus distributing ITNs. ACTs in Vietnam are free in the public sector and there are virtually no private sector sales. With low annual case numbers and a concentration of cases in a few provinces, Vietnam has achieved nearly full ACT coverage for malaria, while implementation of vector control is irregular and generally reaches less than 30% of the population in endemic provinces. This contrasts with the access and treatment scenario in Africa where it was recently reported that only 20% of children under the age of five received an ACT for a confirmed case of *P. falciparum* malaria [29]. As ACT scale-up from 2000 to 2015 reached only 20% coverage in Africa, while ITN scale-up reached coverages between 40% and 70% [30], perhaps it is not surprising that the estimated effect size of ACT on malaria incidence in Africa is low. The differing results in Vietnam and Africa are not necessarily contradictory. It may simply be the case that in Africa we have not yet had an opportunity to fully measure the extent to which ACTs could reduce malaria incidence under a scenario of widespread access to ACTs. In principle, a statistical effect should be detectable for slow increases of ACT coverage, but in practice these effects can be non-linear and an increase from 0% to 20% may not have the same effect as an increase from 20% to 40%. Likewise, in Vietnam, increasing access to ITNs may have reduced incidence even more quickly, but there were no provinces in the data set that showed a rapid scale-up of ITS/IRS coverage. This finding is consistent with descriptions of the control program as it was being implemented in the 1990s [28].

It is also possible that in low transmission areas, ACT case management is more effective than ITN use as a general malaria control policy, whereas the reverse may be true for high transmission areas. ACT case management is a control strategy that works by targeting symptomatic and infected individuals, while vector control acts broadly to protect the entire at-risk population. In addition, ACT case management may be less effective at reducing transmission in highly endemic areas due to the presence of asymptomatic cases that may never be diagnosed and treated [31, 32]. In Vietnam, where transmission intensity is relatively low, population immunity is low and thus individuals with malaria infections are more likely to experience symptoms, seek treatment at a health center, and receive ACT treatment [33, 34]. Additionally, bed nets might have played a less important role in the decline of malaria in Vietnam between 1991-2014 due to the challenge of increasing bed net utilization in remaining high-risk groups such as forest workers who stay overnight in the forest where transmission intensity is the greatest [35]. The relative population-level effects of ACT case management in areas of low transmission may suggest that rapid ACT scale-up could be an effective endgame strategy for regions close to achieving elimination [32].

Urbanization trends in malaria-endemic countries over the past decades have had an effect of reducing the at-risk population for malaria and consequently malaria incidence [36–38]. In central Vietnam, an inverse association between the proportion of the population living in urban areas and the incidence of suspected malaria was observed. However, the uncertainty in the urbanization covariate was large enough to preclude a significant association with confirmed or estimated malaria cases. No associations between incidence and urbanization were found in the southern provinces, and associations were not robust in the north. More detailed spatial data on the distribution of malaria cases in central Vietnam would both contribute to our understanding of urbanization trends and malaria and would help focus control efforts. As the majority (68%) of the Vietnamese population in 2014 still lived in rural areas, it is possible that any effects of urbanization on malaria, if real, would still be too small to detect.

Obtaining a coarse estimate of the positive predictive value (PPV) of malaria clinical diagnosis revealed that the average PPV for the years 2010-2014 in the northern part of the country ranged from 0.10% to 54.44%, whereas in the central and southern parts of the country, PPV ranged from 31.4% to 99.2%. In the northern provinces, suspected case counts are very high, but confirmed case counts are very low. Based on Vietnam’s substantial experience with malaria microscopy and clinical malaria diagnosis, and the existence of a centralized malaria health system, it is likely that over-reporting of suspected malaria cases is occurring in the northern provinces. It is much less likely that a lack of microscopes, microscopists, or a truly low positive predictive value of clinical diagnosis is the cause of the large number of suspected cases reported from the north. Our estimate of malaria incidence is based on (1) the assumption that the true number of cases is greater than the number of confirmed cases and smaller than the number of suspected cases, and (2) that PPV should show little change through time. This is an appropriate way of approximating true malaria incidence in provinces where the suspected and confirmed numbers are close, but it works very poorly in provinces (e.g., many in northern Vietnam) where these two numbers differ substantially.

One general limitation of using suspected malaria cases from national malaria reports is that this quantity will not include true positives that did not report to the health system due to low severity or lack of symptoms. Individuals with mild symptoms (sub-clinical infections) are less likely to seek care and, when they do, are more likely to receive a false negative clinical diagnosis. Asymptomatic and sub-clinical infections are likely to be rare in Vietnam today, where the relatively low transmission intensity reduces the frequency of exposure and the population-level of immunity; however, they cannot be ruled out completely. In highly-endemic areas where asymptomatic cases are more prevalent [39], counts of suspected cases by clinical diagnosis only may grossly underestimate true malaria prevalence.

Two additional data limitations are that the data are aggregated annually and that drug purchase reporting does not allow us to state with certainty which antimalarial treatments were actually used and in what proportions. A long-term cohort or community study with active surveillance and knowledge on antimalarial drug usage, vector control, and other interventions would be the ideal data set for inferring associations between malaria and antimalarial interventions. These study designs are common in Africa where transmission levels are still high [40–44], but the low number of malaria cases in Vietnam would result in a very small number of observed malaria cases unless a highly specific population group were chosen. Retrospectively, only annual data are available and no patient-level drug-use data are available outside of those observed in clinical research studies.

With communicable diseases still playing a large role in the World Health Organization’s health-related Sustainable Development Goals for 2030, malaria elimination will stay on the agenda as an important public health priority in countries that are in or approaching near-elimination phase. Vietnam reported fewer than 10,000 confirmed malaria cases both in 2015 and 2016, placing it in a small group of 30 to 35 countries that could realistically eliminate malaria by 2030 [45]. As monitoring and active surveillance scale up during this phase, it is critical to understand the local causes of malaria decline over the past ten or more years that have enabled each country to reach near-elimination phase. In Vietnam, the path to low malaria incidence has clearly been led by high levels of artemisinin and ACT use in the public sector at coverage rates that can realistically be considered as having a noticeable impact on malaria elimination in some provinces. Active surveillance, reactive case detection, patient compliance, and case follow-up are likely to be key ingredients in Vietnam’s next phase of moving the majority of its provinces to zero malaria over the next decade. The major gloom on the horizon is the spread of artemisinin-resistance genotypes in Vietnam [46, 47], as the arrival of drug-resistance can undermine elimination efforts [48, 49]. It is uncertain if a public health response in this context (e.g., lengthening ACT courses, follow-up with second-line drugs) will be sufficient to maintain the cure rates previously observed with ACTs and keep Vietnam on the road to elimination. Public health agencies and researchers must work together during this time to share knowledge and data, remain open to quick changes in public health strategy, and squarely keep the focus on the public good of eliminating malaria.

## Methods

### Data Collection

The data used in this analysis were obtained from the Institutes for Malariology, Parasitology, and Entomology (IMPE) located in Ho Chi Minh City and the National Institutes for Malariology, Parasitology, and Entomology (NIMPE) in Hanoi. Data structure and cleaning have been described previously [27]. Briefly, NIMPE is responsible for coordinating, funding, and carrying out malaria control activities in Vietnam, including defining guidelines, training staff, supporting local and provincial malaria posts and clinics, purchasing and distributing antimalarial drugs, distributing insecticide-treated nets, identifying malaria transmission hot-spots, and spraying indoor insecticide in homes. NIMPE logs these activities in annual reports summarizing the status of malaria control in each province. The NIMPE reports contain a variety of provincial-level data on malaria cases, vector control measures, antimalarial drug requests, budgetary figures, and other aspects of the control program. The data contained in these reports are collected through a hierarchical reporting system: provincial hospitals and district health centers submit monthly reports to provincial malaria control centers, which then submit provincial summary reports to NIMPE or to one of the regional IMPE offices [50].

Data from these reports were collected in hard copy for the years 1991 to 2014, for all 58 provinces and 5 municipalities in Vietnam, and were input into Microsoft Excel spreadsheets by qualified staff members at the Oxford University Clinical Research Unit in Ho Chi Minh City, Vietnam. In order to account for changes in provincial borders over the study time period, provinces and municipalities were aggregated into 51 administrative units, which we henceforth collectively refer to as ‘provinces’. The following provinces/municipalities were combined for analysis: 1) Bac Can and Thai Nguyen provinces; 2) Bac Ninh and Bac Giang provinces; 3) Bac Lieu and Ca Mau provinces; 4) Binh Duong and Binh Phuoc provinces; 5) Hau Giang, Can Tho, and Soc Trang provinces; 6) Ha Nam and Nam Dinh provinces; 7) Hai Duong and Hung Yen provinces; 8) Quang Nam province and the city of Da Nang; 9) Vinh Phuc and Phu Tho provinces; 10) Dak Lak and Dak Nong provinces; 11) Dien Bien and Lai Chau provinces.

### Malaria Case Data and Case Estimation

Data collected on malaria cases include provincial-level counts of cases diagnosed by clinical observation, cases confirmed by microscopy, cases of *P. falciparum* malaria as confirmed by microscopy, cases of *P. vivax* malaria as confirmed by microscopy, cases of severe malaria, and deaths caused by malaria infection. Data on malaria case numbers were available as either the number of clinically suspected malaria cases, *y,* or the number of suspected cases confirmed by microscopy, *x,* where *x* is a subset of *y* and both may include cases of *P. falciparium* and *P. vivax* malaria. However, for the *y* – *x* cases that remained unconfirmed, the reports do not indicate whether these cases tested negative by microscopy or if they were simply untested. The distinction between cases that tested negative and untested cases could be made if either the positive predictive value, *q,* of clinical diagnosis or the fraction of clinically suspected cases that undergo blood-slide diagnosis, *f_BSD_*, were known. If *f_BSD_* is known, then the total number of suspected cases that undergo microscopy, *y* · *f_BSD_*, can be used to calculate the positive predictive value, *q,* where *q* has a maximum value of one:

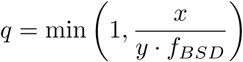

Likewise, if the positive predictive value is known, then the fraction of suspected cases that underwent blood-slide confirmation can be calculated, unless *q* = 1, in which case all clinically suspected malaria cases are truly malaria. If the rate of blood-slide confirmation, *f_BSD_*, is known, then the true number of malaria cases (*falciparium* and *vivax*) can be estimated as:

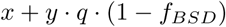

which is equal to 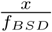 when *q* ≤ 1. Since independent estimates of *q* or *f_BSD_* cannot be obtained, we assumed that *f_BSD_* increased linearly in each province during the years 1991-2014. This assumption is consistent with the NIMPE annual reports, which show an increase in health system capacity and microscopy during this period. Assuming a linear increase in *f_BSD_* yields 24 individual values of *q* for the years 1991-2014. The linear increase in *f_BSD_* is chosen to minimize the variance in the year-to-year positive predictive value, as it is likely that clinicians’ ability to correctly diagnose a malaria case did not change significantly during this time. This assumption is supported by the opinions of IMPE/NIMPE staff. However, the relative prevalence of other febrile diseases is likely to affect the positive predictive value of malaria clinical diagnosis, depending on the level of similarity between the symptoms of these diseases and malaria symptoms. Unfortunately, incidence data for other febrile diseases are not available.

### Predictor Data

We identified and collected provincial-level data on potential predictors of provincial malaria case counts in Vietnam from 1991-2014. We used data on vector control measures contained in the annual reports to calculate the proportion of the population protected by vector control measures, as has been done in previous work [27]. These data include the total number of people protected by insecticide-treated bed nets (ITNs) and indoor residual spraying (IRS), reported as the sum of these two measures. ITN coverage is defined by NIMPE as the fraction of households possessing an ITN that was shared by no more than three other individuals[51]. Imputation by linear interpolation was used to impute missing data on vector control measures.

We also looked at provincial-level data on the proportion of drug purchases for *P. falciparum* malaria that contained artemisinin. Data on the use of antimalarial drugs were contained in these reports in the form of requests placed by provincial malaria control programs to NIMPE for antimalarial drugs based on projected need for the subsequent year. Requests for antimalarial drugs likely capture true antimalarial drug usage in the country because all public hospitals and health centers in Vietnam provide antimalarial drugs free of charge to individuals suspected of having malaria [27, 50]. We looked at treatments for *P. falciparium* malaria only because the vast majority of suspected malaria cases were not confirmed by blood-smear for the first half of the time series, and unconfirmed cases were typically presumed to be *falciparium* and treated as such. Thus, in the analysis, we investigated the association between malaria incidence *(falciparium* and *vivax* combined) and the degree to which artemisinin-based drugs were the dominant purchases by local malaria control programs. In a setting where all suspected malaria is confirmed as *P. falciparium* or *P. vivax,* artemisinin would only act to reduce falciparum transmission; however, when the majority of cases are unconfirmed, the presumption of falciparum and treatment with artemisinins will act to reduce onward transmission of both falciparum and vivax.

Drug purchase data were available for all but three years across the study period (1991, 1993, and 1996) and were specified in terms of the drug type and number of units requested (e.g., tablets, pills, tubes). Appendix Table S1 shows the antimalarial drugs ordered by province according to the reports. Since a complete course of treatment for an individual patient usually requires multiple drug units (doses), the number of drug units ordered was converted to the number of courses of treatment for *P. falciparum* malaria ordered using dose-to-course conversions. These conversions were taken from the national treatment guidelines contained in the annual reports and obtained from conversations with IMPE staff (See Appendix Table S2). For drugs that were used to treat more than one type of parasite, the number of courses of treatment for *P. falciparum* was calculated by dividing the number of drug units ordered by the average number of drug units per treatment course and multiplying by the fraction of cases that were *P. falciparum* across all provinces in the corresponding year. The average number of drug units per treatment was calculated as the weighted average of the recommended dosages, using the year-specific relative proportions of *P. falciparum* cases and *P. vivax* cases across all provinces as weights. In order to calculate the proportion of treatment courses that contained artemisinin, the number of treatment courses for *P. falciparum* was then summed for the drugs that contained artemisinin and divided by the number of treatment courses for *P. falciparum* summed across all drug types. As there was uncertainty about whether primaquine was always used in conjunction with an ACT or whether it was sometimes used alone, an alternative dose-to-course conversion was considered (see Appendix Figure S2). Where possible, simple linear averaging was used to impute missing data on antimalarial drug requests. For the year 1991, antimalarial drug requests were imputed by linearly extending the 1992 and 1993 trend back to 1991 and verifying that the imputed value was less than the values in 1992 and 1993.

The reports also contained data on two measures of health system capacity: the discretionary budget and the training of health care workers. Budgetary data, broken down into sub-budget allocations for the local malaria control program, and data on staff trained to support the malaria control program, were both available in the reports by province and year. We used the discretionary budget per capita, which excludes the allocated budget for purchasing antimalarial drugs, insecticides, and subsidies for ITNs and IRS, as a measure of health system capacity after adjusting for historic inflation in the VND. We also used the number of staff trained per capita as a measure of health system capacity. Data for these two measures of health system capacity were only available for the years 1997-2014. Due to the high degree of missingness in measures of health system capacity, no imputations were conducted.

Data on the number of individuals living in urban areas were available by province for all years after 1994 from the General Statistics Office of Viet Nam and are available online at the GSO website [52]. These data were used to calculate the proportion of the population in each province living in urban areas. Total population data was also available for all years after 1993 from the General Statistics Office of Viet Nam. Missing population data were imputed linearly.

### Statistical Analysis

We fit Poisson-regression models to our provincial-level malaria case data from 1991-2014. Three measures of malaria cases were separately used as model outcomes: 1) clinically-diagnosed malaria cases (i.e., ‘suspected’ cases), 2) blood-smear confirmed cases (i.e., ‘confirmed’ cases), and 3) the estimated true number of malaria cases (i.e., ‘estimated’ cases) which we obtained by minimizing the year-to-year variation in the positive predictive value of clinical diagnosis, as described above. All case measures include cases of both *falciparium* and *vivax* malaria. Five variables, measured by province and by year, were identified as potential predictors of incidence: 1) the proportion of treatment courses for *P. falciparum* containing artemisinin, 2) the proportion of the population protected by vector control measures, 3) the proportion of the population living in urban areas, 4) the discretionary budget per capita for the malaria control program, and 5) staff trained per 100 persons. Our models included a province-specific fixed effect and the log of the population as an offset term. We used Generalized Estimating Equations (GEE), appropriate for correlated time-series data, to fit our models, and specified a log-link function, an independent correlation structure, and a robust covariance matrix estimator [53]. We also used non-parametric Spearman’s rank correlation tests to investigate temporal trends in potential predictors of case counts.

We first fit the models to data from all 51 provinces in Vietnam for the 24 years from 1991-2014. Then, we stratified our province-level data into northern, central, and southern regions and separately fit the models in order to explore whether the relationship between malaria incidence and malaria control measures varied regionally. Regional groupings of provinces are shown in Appendix Figure S1.

Due to missingness in the health system capacity variables, we fit two types of models: models using all five covariates as predictors (i.e., ‘five-covariate models’) and models using only three covariates (i.e., ‘three-covariate models’ excluding the two measures of health system capacity – discretionary budget per capita and staff trained per 100 persons. The three-covariate models included data from 1991-2014, whereas the five-covariate models included data from 1997-2014. All analysis was conducted in R, using the *geepack* package for fitting generalized estimating equations [54]. Shape files for creating maps of Vietnam were obtained from the GADM database of Global Administrative Areas.

## Acknowledgments

SMG was supported by a grant from the Princeton University Council for International Teaching and Research. PDT, DDHG, NVT, TDN, and TTH are supported by Wellcome Trust grant 089276/B/09/7. MFB was supported by a Wellcome Trust/Royal Society Sir Henry Dale Fellowship (098511/Z/12/Z) and is currently supported by the Pennsylvania State University.

### Disclosures

JF is the director of the Wellcome Trust. The authors declare that they have no other conflicts.

## Appendix

**Figure S1.**
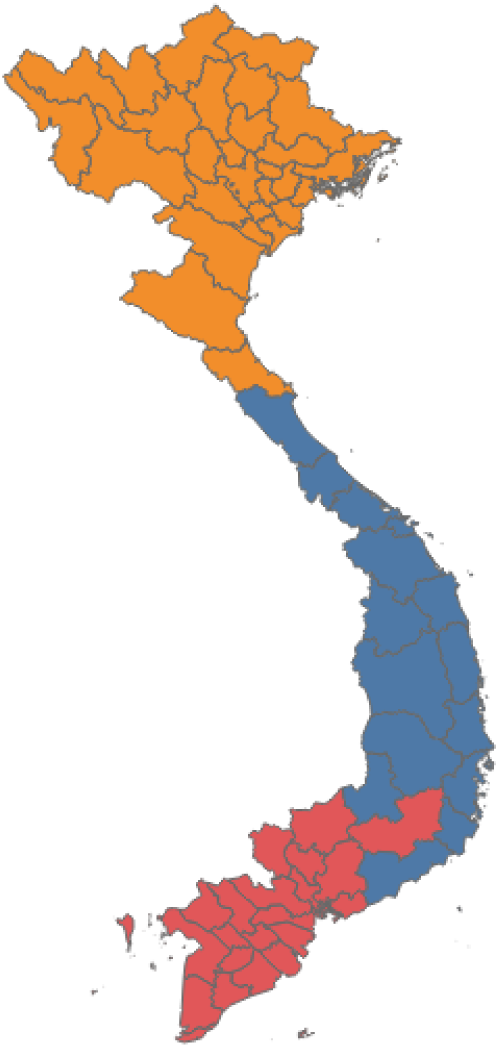
Northern (orange), central (blue), and southern (red) regional grouping of provinces used in the analysis.

**Figure S2.**
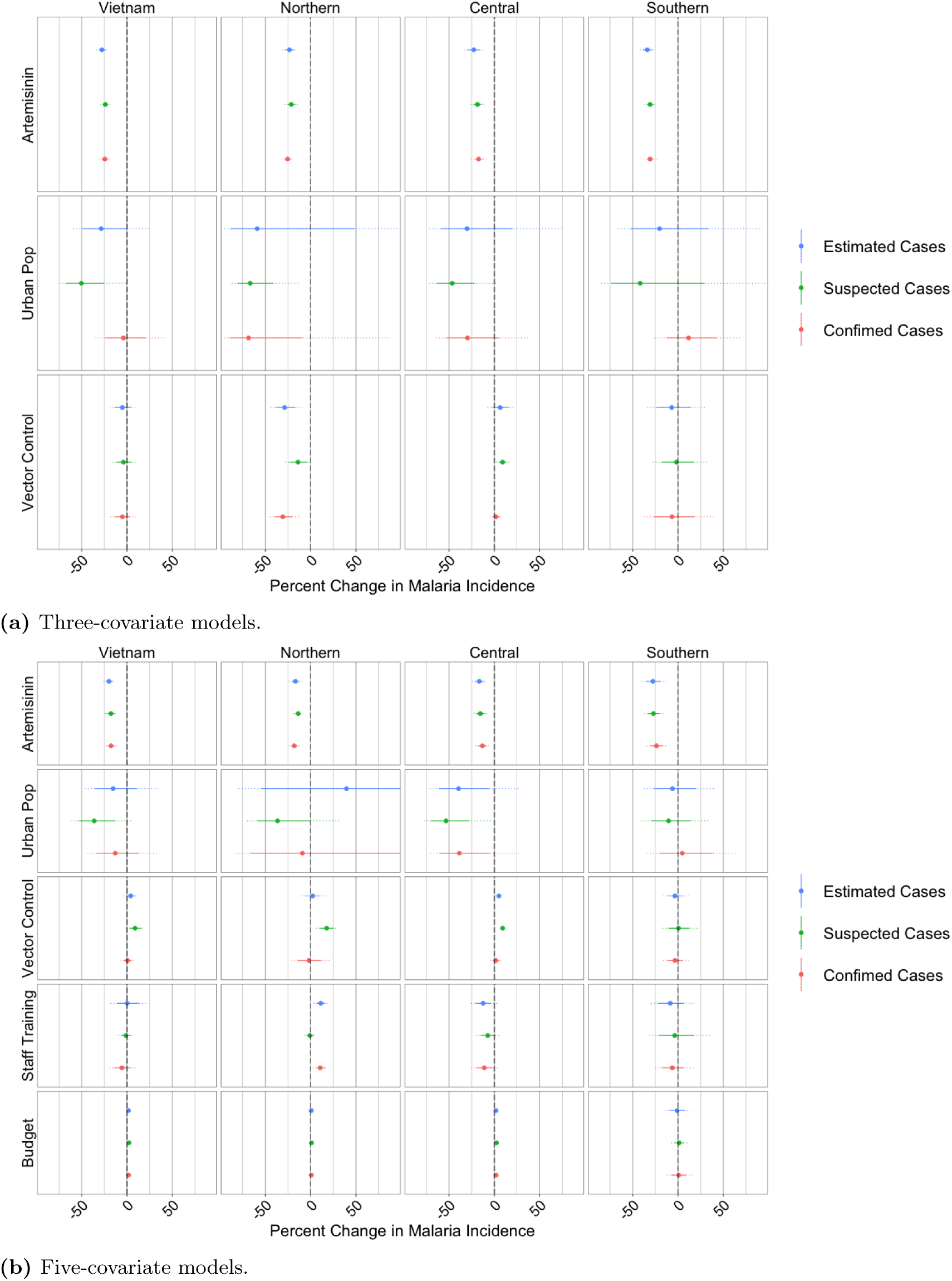
Regression results using the alternative calculation for the proportion of treatments containing artemisinin, shown as the percent change in incidence associated with a 10% increase in the listed covariate, by region and for all provinces in Vietnam for a) the models including three covariates only, and for b) the models that included two additional covariates as measures of health system capacity. The circle shows the mean effect size, the solid line shows the 95% confidence interval, and the dotted lines shows the 99.9% confidence interval. Model outcome (estimated, suspected, or confirmed cases) is indicated by effect size color. For clarity, the x-axis has been limited to range from -90 to 90.

**Table S1.** Anti-malarial drugs ordered by province, 1991-2014

| Drug | Unit Weight | Years Requested |
| --- | --- | --- |
| DHA-PPQ | 40 mg DHA/ 320 mg Piperaquine | 2007-2014 |
| CV8 | 32 mg DHA/ 320mg PPQ/ 90 mg trimethoprim/ 5 mg primaquine | 2001-2006 |
| Artemisinin | 250 mg | 1992-2003 |
| Artesunate | 50 mg | 1992-2008 |
| Artesunate Injection | 60 mg | 1994-2014 |
| Quinine | 250 mg | 1992-2001; 2004-2005; 2007-2008; 2011-2014 |
| Quinine Injection | 500 mg | 1992-2001; 2004-2005 |
| Quinoserum | 50 mg | 1992-1995 |
| Primaquine | 13.2 mg | 1992-2014 |
| Chloroquine | 250 mg | 1992-2014 |
| Mefloquine | 250 mg | 1994-1997; 1999- 2002 |
| SR2 | 500 mg Sulfadoxine/25 mg Pyrimethamine | 1992-2003 |
| SR3 | probably Sulfadoxine / Pyremethamine / Piperaquine (no guideline from MoH) | 1992 |
| Fansidar Injection | 400mg Sulfadoxine/20mg Pyrimethamine 2.0 mL | 1995 |

*Note.* DHA = Dihydroartemisinin

**Table S2.** Malaria Treatment Guidelines

| 2009-2014 Guidelines | Drug (Avg. units per treatment course) |
| --- | --- |
| Uncomplicated *P. falciparum* | DHA-Piperaquine (8 tablets) + Primaquine (4 tablets) |
| Uncomplicated *P. falciparum*<br>(in 1st trimester pregnancy) | Quinine (42 tablets) + Clindamycin (14 tablets) |
| Severe *P. falciparum* | Artesunate Injection (4-10 tubes with switch to oral) |
| *P. vivax* | Chloroquine (10 tablets) + Primaquine (28 tablets) |
| 2003-2008 Guidelines |  |
| Uncomplicated *P. falciparum* | Artesunate (16 tablets) + Primaquine (4 tablets)<br>OR CV8 (8 tablets) OR DHA-PPQ (8 tablets) |
| Uncomplicated *P. falciparum*<br>(in 1st trimester pregnancy) | Quinine (42 tablets) |
| Severe *P. falciparum* | Artesunate Injection (2-9 tubes with switch to oral) |
| *P. vivax* | Chloroquine (10 tablets) + Primaquine (20 tablets) |
| 1991-2002 Guidelines |  |
| Uncomplicated *P. falciparum* | Chloroquine (8 tablets) + Primaquine (4 tablets) |
| Uncomplicated *P. falciparum*<br>(with chloroquine resistance) | Fansidar (3 tubes)<br>OR Fansidar (3 tubes) + Quinine (42 tablets)<br>OR Quinine (42 tablets) + Tetracycline (28 tablets)<br>OR Mefloquine (3 tablets)<br>OR Artemisinin (16 tablets, 7 days) + Primaquine (4 tablets) |
| Severe *P. falciparum* | Artesunate Injection (2-9 tubes)<br>OR Quinine Injection (3 tubes/day for 7 days) |
| *P. vivax* | Chloroquine (8 tablets) + Primaquine (20 tablets) |

